# Red Sea SAR11 and *Prochlorococcus* Single-cell Genomes Reflect Globally Distributed Pangenomes

**DOI:** 10.1101/549816

**Authors:** Luke R. Thompson, Mohamed F. Haroon, Ahmed A. Shibl, Matt J. Cahill, David K. Ngugi, Gareth J. Williams, James T. Morton, Rob Knight, Kelly D. Goodwin, Ulrich Stingl

**Affiliations:** Red Sea Research Center, King Abdullah University of Science and Technology, Thuwal, Saudi Arabia; Department of Biological Sciences and Northern Gulf Institute, University of Southern Mississippi, Hattiesburg, MS, United States of America; Ocean Chemistry and Ecosystems Division, Atlantic Oceanographic and Meteorological Laboratory, National Oceanic and Atmospheric Administration, stationed at Southwest Fisheries Science Center, La Jolla, CA, United States of America; School of Ocean Sciences, Bangor University, Anglesey, United Kingdom; Department of Pediatrics, University of California San Diego, La Jolla, CA, United States of America; Department of Computer Science and Engineering, University of California San Diego, La Jolla, CA, United States of America; Center for Microbiome Innovation, University of California, San Diego, CA, United States of America; Department of Microbiology & Cell Science, Fort Lauderdale Research and Education Center, UF/Institute of Food and Agricultural Sciences, University of FL, Davie, Florida, United States of America

**Keywords:** metagenomics, *Pelagibacter*, population genomics, SAG, single-cell genomics

## Abstract

Evidence suggests many marine bacteria are cosmopolitan, with widespread but sparse strains poised to seed abundant populations upon conducive growth conditions. However, studies supporting this “microbial seed bank” hypothesis have analyzed taxonomic marker genes rather than whole genomes/metagenomes, leaving open the possibility that disparate ocean regions harbor endemic gene content. The Red Sea is isolated geographically from the rest of the ocean and has a combination of high irradiance, high temperature, and high salinity that is unique among the ocean; we therefore asked whether it harbors endemic gene content. We sequenced and assembled single-cell genomes of 21 SAR11 (subclades Ia, Ib, Id, II) and 5 *Prochlorococcus* (ecotype HLII) cells from the Red Sea and combined them with globally-sourced reference genomes to cluster genes into ortholog groups (OGs). Ordination of OG composition could distinguish clades, including phylogenetically cryptic *Prochlorococcus* ecotypes LLII and LLIII. Compared with reference genomes, 1% of *Prochlorococcus* and 17% of SAR11 OGs were unique to the Red Sea genomes (RS-OGs). Most (83%) RS-OGs had no annotated function, but 65% of RS-OGs were expressed in diel Red Sea metatranscriptomes, suggesting they could be functional. Searching *Tara* Oceans metagenomes, RS-OGs were as likely to be found as non-RS-OGs; nevertheless, Red Sea and other warm samples could be distinguished from cooler samples using the relative abundances of OGs. The results suggest that the prevalence of OGs in these surface ocean bacteria is largely cosmopolitan, with differences in population metagenomes manifested by differences in relative abundance rather than complete presence-absence of OGs.

## Importance

Studies have shown that as we sequence seawater from a selected environment deeper and deeper, we approach finding every bacterial taxon known for the ocean as a whole. However, such studies have focused on taxonomic marker genes rather than on whole genomes, raising the possibility that the lack of endemism results from the method of investigation. We took a geographically isolated water body, the Red Sea, and sequenced single cells from it. We compared those single-cell genomes to available genomes from around the ocean, and to ocean-spanning metagenomes. We showed that gene ortholog groups found in Red Sea genomes but not in other genomes are nevertheless common across global ocean metagenomes. These results suggest that Baas Becking’s hypothesis “everything is everywhere, but the environment selects” also applies to gene ortholog groups. This widely dispersed functional diversity may give oceanic microbial communities the functional capacity to respond rapidly to changing conditions.

## Introduction

Marine bacteria thrive throughout the surface ocean despite low nutrients, high irradiation, and other physicochemical stressors. Adaptations enabling survival can be at the level of transcriptional, translational, and other methods of cellular regulation that occur at time-scales of minutes to hours (1, 2). Alternatively, microbial genomes can evolve new functions on the scale of thousands to millions of generations (3, 4). Evolution via horizontal gene transfer enables the introduction of entirely new functionality (gene gain) as well as genome streamlining (gene loss) for more efficient resource (e.g., nitrogen, phosphorus) allocation (5). Therefore, it is expected that the genomes of marine bacteria will display differences in gene content correlated with the physicochemical environment in which they live. Indeed, both individual genomes (cultured and single-cell genomes) (6–10) and community genomes (metagenomes) (11, 12) show that bacteria in the oligotrophic (nutrient-poor) surface ocean carry streamlined genomes finely tuned to their environments.

Examples of adaptive gene presence-absence patterns are seen in the most numerous groups of bacteria in the oligotrophic tropical and sub-tropical surface ocean, the photoautotrophic picocyanobacteria *Prochlorococcus* and *Synechococcus* and the chemoheterotrophic Alphaproteobacteria SAR11 clade (*Candidatus* Pelagibacter ubique). Genomes of these genera are smaller than their relatives in less nutrient-poor environments (6, 8), suggestive of genome streamlining to conserve resources used for genome replication (3). Consistent with genome streamlining, the genes maintained in *Prochlorococcus* and SAR11 genomes are correlated with physical features in parts of the water column in which they are found, for example, genes for acquisition of nitrite and nitrate in genomes found where those compounds are available (3, 8). Examples revealed through comparative community genomics include an enrichment of phosphorus acquisition gene ortholog groups in the Atlantic relative to the Pacific Ocean (11, 13) and an enrichment in osmolyte oxidation gene ortholog groups in the Mediterranean and Red Seas relative to the Atlantic and Pacific Oceans (12).

The Red Sea is an attractive environment for the study of genomic adaptations. Geographically, the Red Sea is largely isolated from the rest of the World Ocean, with only a small sill (the Bab el Mandeb) connecting it to the Indian Ocean (14). Among surface waters catalogued in the World Ocean Database, the Red Sea lies at the high end of the global temperature distribution and is higher than any other sea in the global salinity distribution (Fig. S1). The Red Sea, straddling the Tropic of Cancer, experiences year-round high irradiance, and cloud cover across North Africa and the Arabian Peninsula is among the lowest on the planet (NASA Aqua satellite MODIS sensor). The Red Sea is also oligotrophic, with production thought to be limited by nitrogen (15).

Evidence of genomic adaptation to high light and high salinity in the Red Sea has been revealed through comparative metagenomics, showing increased relative abundance of known gene ortholog groups in *Prochlorococcus* and SAR11 (12). Relative to the North Pacific, Sargasso Sea, and Mediterranean Sea, the Red Sea *Prochlorococcus* population had increased frequencies of high-light stress and DNA repair gene ortholog groups (12), the latter likely an adaptation to UV-induced DNA damage. Relative to these same seas, the SAR11 population had increased frequencies of gene ortholog groups for osmolyte degradation (12); osmolytes are important molecules for surviving high salinity in many organisms. Across 45 metagenomes along latitudinal and depth gradients from the surface to 500 m in the Red Sea, temperature explained more variation in gene ortholog groups than any other environmental parameter, and the relative abundance of gene ortholog groups linked to high irradiance, high salinity, and low nutrients were correlated with those parameters (16).

The above-mentioned patterns observed in comparative metagenomics studies were all based on relative abundance of known gene ortholog groups, dependent on a reference genome database with no representatives from the Red Sea. Therefore, the question remains if there are gene functions in the *Prochlorococcus* and SAR11 populations in the Red Sea not found in any other *Prochlorococcus* and SAR11 populations in the ocean. Because of its relative geographic isolation, we might expect the Red Sea to be genetically isolated, with endemic genomic adaptations to its unique combination of high solar irradiance, high temperature, high salinity, and low nutrient levels. Newly identified gene ortholog groups could be informative for understanding microbial adaptation and mechanisms of stress tolerance, and have potential biotechnological applications.

The question of whether there are genetic functions found in only one sea of the global ocean speaks to theoretical questions of microbial biogeography as well. A prevailing idea in microbial ecology is that most microbial species are found at a given site provided the conditions are conducive for their growth. This is known as the Baas Becking hypothesis: “Everything is everywhere, but the environment selects” (17). Among microbial taxa found in seawater, there is growing evidence for a cosmopolitan distribution of these taxa throughout the global ocean (18, 19). Support for the “microbial seed bank” hypothesis has come from deep sequencing of ocean samples, revealing for example that nearly all 16S rRNA operational taxonomic units (OTUs) from a deep-sea hydrothermal vent can be found in the open ocean (19), and that we approach identifying all OTUs in the ocean as sequencing effort increases for a single marine sample (18). Despite this evidence supporting a cosmopolitan distribution of OTUs throughout the ocean, these amplicon sequences (16S rRNA OTUs) are only taxonomic proxies and do not represent the extensive gene-level diversity in microbial genomes. Even if such marker gene sequences are omnipresent across the ocean, genome evolution and diversification, e.g., via horizontal gene transfer, could be occurring that generates gene-level adaptations that are endemic to particular locations. Are microbial gene ortholog groups, defined at the level of genus (SAR11 or *Prochlorococcus*), as widely distributed as microbial 16S rRNA gene sequences?

Here, to investigate microbial genomic diversity in SAR11 and *Prochlorococcus*, including possible endemic adaptation in Red Sea populations, we have sequenced single-cell amplified genomes (SAGs) from the Red Sea and compared their gene ortholog group (OG) content to genomes and metagenomes from around the World Ocean. We have quantified expression of OGs in metatranscriptomes from the Red Sea collected over two consecutive 24-hour day–night cycles. This effort has resulted in 21 SAR11 SAGs, including the first genomes from subclades Ib and Id, and 5 *Prochlorococcus* SAGs. Using these Red Sea SAGs and the OGs they contain as queries for genomic and metagenomic analyses, we have analyzed globally-sourced genomes and metagenomes to investigate the extent to which OGs from surface-ocean *Prochlorococcus* and SAR11 are distributed across the World Ocean.

## Materials and Methods

### Sample collection

A single seawater sample (100 mL) was collected in a polycarbonate bottle from the surface (depth of 0 m) of an open-ocean site in the east-central Red Sea (19.75 °N, 40.05 °E), near the Farasan Banks region, on June 15, 2010. The sample was preserved with dimethyl sulfoxide (5% final concentration), flash frozen in liquid nitrogen, and stored at −80 °C.

Seawater samples for metatranscriptomics were taken March 3-5, 2013, from an open-ocean site in the Red Sea (Kebrit Deep, 24.7244 °N, 36.2785 °E). One sample per depth was collected every 4 h over a 48-h period at four depths: surface (10 m), below the mixed layer (40 m; bottom of mixed layer was 35 m), chlorophyll maximum (75 m), and oxygen minimum zone (420 m). For each timepoint and depth, 1 L seawater was filtered using a peristaltic pump with two in-line filters in series: a 1.6-μm GF/A pre-filter (Whatman), then a 0.22-μm Sterivex filter (Millipore). RNAlater (QIAGEN) was added immediately to fill the dead space of the Sterivex filter, which was then flash frozen in liquid nitrogen and stored at −80 °C.

### Nucleic acid extraction and amplification

Single bacterioplankton cells in the preserved samples were flow-sorted, whole-genome amplified (MDA, multiple displacement amplification), and PCR-screened at the Bigelow Laboratory Single Cell Genomics Center (SCGC, Boothbay Harbor, ME, USA), following previously described protocols (20), with SYTO-13 nucleic acid stain used to stain cells for flow-sorting. SAG identification was carried out with SCGC protocol S-102 for bacteria using 16S rRNA primers 27F and 907R (21, 22). A total of 21 and 5 cells were identified from 16S PCR screening and subjected to a second round of MDA before sequencing. The 16S rRNA gene sequences are available from the European Nucleotide Archive with accession numbers LN850141–LN850161.

The RNA extraction protocol for metatranscriptomics was adapted from (23–25). After expelling RNAlater from the Sterivex filter, 2 mL lysozyme solution (1 mg/mL in lysis buffer: 40 mM EDTA, 50 mM Tris pH 8.3, 0.73 M sucrose) was added, then filter incubated at 37 °C with rotation for 45 min. Proteinase K solution (50 μL at 20 mg/mL, QIAGEN/5PRIME) and SDS solution (100 μL at 20%) were added, then filter incubated at 55 °C with rotation for 2 h. Lysate was expelled to a separate tube; meanwhile, 1 mL lysis buffer was added to the filter to wash at 55 °C for 15 min. The two lysates were pooled, to which was added 1.5 mL absolute ethanol. RNA was then extracted from this solution using the RNeasy Protect Bacteria Midi Kit (QIAGEN). RNA was eluted with two volumes of RNase-free water. RNA sample was concentrated using a speed vacuum, from 250 μL to 60 μL. To this volume we added DNase (1 μL Ambion TURBO DNA-free, 6 μL 10x buffer, 60 μL RNA) and incubated at 37 °C for 30 min. This solution was purified using the RNeasy MinElute Cleanup Kit (QIAGEN) and eluted with RNase-free water. Final yield was 1-2 ng total RNA. Total RNA was amplified using the C&E Version ExpressArt Bacterial mRNA Amplification Nano Kit, which preferentially amplifies mRNA (independent of poly-A tail) and selects against rRNAs. A single round of amplification was performed on 2–4 ng of total RNA which yielded about 10 μg final amplified RNA.

### Nucleic acid sequencing

For single-cell genome sequencing, genomic library preparation with Illumina TruSeq and sequencing with Illumina GAIIx and Illumina HiSeq 2000 was done at the KAUST Bioscience Core Laboratory, generating paired 105-bp reads. The assembled contigs (assembly methods below) are available from NCBI with accession numbers PRJEB9287 (BioProject) and SAMEA3368552–SAMEA3368577 (BioSample), and can also be visualized in Integrated Microbial Genomes system (26) under accession numbers 2630968236, 2630968238–630968254, 2630968277–2630968281, and 2630968285–2630968287.

For metatranscriptomics, sequence data were processed as described in (27). Amplified RNA was used to construct sequencing libraries using the TruSeq Stranded RNA LT Sample Prep Kit (Illumina) according to the manufacturer’s protocol. Libraries were paired-end sequenced with the Illumina HiSeq 2000 platform (2 × 100 bp). Raw RNA sequences have been deposited in NCBI GenBank with Bioproject number PRJNA289956. Low-quality reads and sequencing adapters were removed using Trimmomatic v0.32 (28). Sequence reads shorter than 50 bp were discarded. Bowtie 2 v2.2.4 (29) was used to identify and remove PhiX contamination sequences. The remaining sequences were error-corrected using the BayesHammer algorithm (30) implemented in the SPAdes v3.5.0 (31), followed by removal of putative ribosomal RNA (rRNA) gene transcripts with SortMeRNA v2.0 (32).

### Genome assembly and annotation

De novo assemblies were generated using CLC Genomics Workbench 4.9. The genomes were assembled independently and, unless otherwise specified, the following applies to all of the SAGs. The reads were first imported and quality trimmed with a limit of 0.01. They were then assembled using CLC’s *de novo* assembler with a word size (*k*-mer) of 64 and with the min/max of the insert size set to 100/1000 bp. Only those contigs greater than 200 bp in length were included in downstream analyses. The reads were mapped to the consensus sequence of the assembled contigs using CLC’s default parameters but with the length fraction set to 1.0 and the similarity set to 0.95.

Assembled SAG contigs were ordered and oriented relative to SAR11 HTCC1062 (NC_007205.1) or *Prochlorococcus* MIT 9202 (NZ_DS999537) using ABACAS 1.3.1 (33). The ordered sequences were then imported into GAP4 (34) and additional joins were made between overlapping contigs if conserved synteny supported the arrangement. To identify and remove possible contaminating sequences from the assemblies, each contig was retained only if it met one or both of the following criteria: (i) the contig was binned into a bin annotated as SAR11 or *Prochlorococcus* using Metawatt 3.5 (35), using the “medium” bin level, with a minimum bin size of 50 kbp and minimum contig size of 500 bp; (ii) the contig had a top-10 BLASTN hit against GenBank nt, with E-value <1e–5, to SAR11 or *Prochlorococcus*.

Prediction of gene open reading frames (ORFs) and functional annotation of SAGs was performed by the RAST web service (36) with FIGfam Release 59.

### Ortholog group clustering

Predicted proteins from SAGs were clustered with proteins from published cultured and SAG genomes (supplemental file 1) into ortholog groups (OGs) using OrthoMCL 2.0 (37). OrthoMCL configuration settings were as follows: percentMatchCutoff=50, evalueExponentCutoff=−5. This yielded 5272 SAR11 OGs and 10439 *Prochlorococcus* OGs. After OrthoMCL clustering, OGs were assigned as core and non-core based on copy number in the non-Red Sea, cultured (non-SAG) genomes: core OGs are those found at least once in each of the non-Red Sea, cultured genomes, and non-core OGs are those not found in at least one of the non-Red Sea, cultured genomes. Among SAR11, there were 683 core OGs and 4589 non-core OGs. Among *Prochlorococcus*, there were 1152 core OGs and 9287 non-core OGs.

### Estimation of genome completeness

Completeness of SAGs was assessed using two methods. First, completeness was assessed using single-copy ‘core’ OGs, i.e., those OGs found once and only once in each complete genome based on the OrthoMCL clusters (analyzed separately for SAR11 and *Prochlorococcus*). Completeness was calculated as the number of core orthologs present in each SAG out of 649 SAR11 or 1144 *Prochlorococcus* single-copy core OGs. Second, genome completeness of the SAGs was assessed using CheckM 1.0.3 (38) using the lineage-specific workflow (lineage_wf). CheckM was also used to estimate genome redundancy (called “contamination” in CheckM).

### Genome taxonomy and phylogenetics

A total of 89 SAR11 and 96 *Prochlorococcus* shared single-copy orthologous genes were identified using the GET_HOMOLOGUES software (39). Amino acid sequences translated from gene sequences were aligned using the MAFFT software (40). These alignments were concatenated, sites with gaps were deleted, and the concatenated data were partitioned using the PartitionFinder software (41) to account for variations of evolutionary processes among gene families. With the Bayesian information criterion (BIC) statistic, a 16-partition framework was chosen to optimally describe the variability, in which the LG rate matrix with Gamma distribution of rate variation (LG+G) was selected for 15 partitions and the VT rate matrix with Gamma distribution of rate variation (VT+G) was selected for the remaining partition. This partition model was used in the maximum-likelihood phylogenomic construction using the RAxML software (42).

### Ordination of SAGs and genomes using *k*-mer composition and ortholog composition

SAGs and reference genomes (Table S1) were analyzed using principal components analysis (PCA) of nucleotide composition and OG composition. Nucleotide composition of the SAGs and reference genomes (SAR11 and *Prochlorococcus* scaffolds >200 kbp from Integrated Microbial Genomes, https://img.jgi.doe.gov) was determined as 6-nucleotide words or *k*-mers (6-mers). *k*-mer frequencies were calculated using Jellyfish 2.2.5; the main command used was jellyfish count −m 6 −t 8 −s 1M. This resulted in a table of 6-mer frequencies in the SAGs and genomes, one table each for SAR11 and *Prochlorococcus*. OG composition was derived from tables of OrthoMCL clusters, which were subsampled so that all genomes had the same number of gene counts in the table: the OG composition tables (with counts of 5272 unique SAR11 OGs and 10439 unique *Prochlorococcus* OGs) were subsampled down to 800 gene counts per SAR11 SAG (keeping 12 of 21 SAGs) and 1400 gene counts per *Prochlorococcus* genome (keeping 5 of 5 SAGs). Prior to PCA, a pseudo-count of 1 was added to *k*-mer and OG count tables to account for zero values; *k*-mer counts were then converted to relative abundances for each genome (unnecessary for OG counts because of the subsampling procedure); *k*-mer relative abundances were then standardized to *z*-scores (not done for OG counts because this reduced the resolving power of PCA). PCA was then performed using the Scikit-Learn function sklearn.decomposition.PCA (43).

### Mapping of metatranscriptomic reads to OGs

The quality-filtered mRNA reads from the 52 samples were mapped against the SAGs using Bowtie 2 (29) with default settings. Each read mapping above the threshold was assigned to exactly one gene in a SAG contig. The resultant read counts were normalized based on the FPKM metric (fragments per kilobase of gene per million mapped reads). Per-sample FPKM counts for each gene were then summed by OGs, resulting in per-sample FPKM counts for each OG. For downstream analysis, counts were converted to a simple presence–absence measure: if any gene belonging to the OG had one or more mapped transcript, that OG was marked as present in that sample.

### Detection and rarefaction analysis of OGs in Tara Oceans metagenomes

A set of 139 prokaryote-enriched *Tara* Oceans metagenomic gene files (44) was downloaded from the European Nucleotide Archive (https://www.ebi.ac.uk/ena, ERZ096909–ERZ097150). Each file contains nucleotide sequences for genes predicted on *Tara* Oceans metagenomic contigs that were assembled from shotgun sequencing reads from individual *Tara* Oceans samples. The prokaryote fraction was 0.22–1.6 μm for stations 004–052 and 0.22–3 μm for stations 056–152; the environmental features of the samples were indicated as “SRF” (surface), “MIX” (mixed layer), “DCM” (deep chlorophyll maximum), and “MES” (mesopelagic zone). The metagenomic gene sequences were queried against a database of translated proteins from the SAGs and genomes with DIAMOND 0.8.26 (45) using the program blastx with parameters –p 40–k 25 –e 1e–3. The top hit (SAG or genome protein sequence) for each *Tara* gene sequence (E-value < 1e–5) was retained. E-value cutoffs of 1e–10 and 1e–15 were also tested, which showed the same trends as E-value < 1e–5 but with fewer total OGs identified. Counts of the number of times each protein was a top hit were then summed across each OG. This resulted in a table of OGs by samples where each OG was either present (at least one constituent protein was a top hit at least once) or absent in each sample. These presence–absence tables (one for SAR11, one for *Prochlorococcus*) were used to generate rarefaction curves: samples were added one-by-one randomly (1000 permutations), and the cumulative number of OGs found was recorded.

### Ordination of *Tara* Oceans metagenomes by OG composition

OG counts (total, not presence–absence) in *Tara* Oceans surface (SRF) sample metagenomes were used for ordination using PCA. Prior to PCA, a pseudo-count of 1 was added to OG count tables to account for zero values; counts were then converted to relative abundances for each metagenome; OGs with an average relative abundance across all metagenomes less than 0.0001 (0.01%) were removed; relative abundances were then standardized to *z*-scores. PCA was then performed using the Scikit-Learn function sklearn.decomposition.PCA (43).

### World Ocean temperature and salinity data

Surface temperature and salinity data (WOD13_ALL_SUR_OBS) from the World Ocean Database 2013 (https://www.nodc.noaa.gov/OC5/WOD13/) were downloaded from the Research Data Archive at the National Center for Atmospheric Research (https://rda.ucar.edu/datasets/ds285.0/).

## Results and Discussion

### Single-cell genome properties and taxonomic classification

Following collection of surface seawater from the east-central Red Sea, flow sorting, and amplification, we sequenced and assembled 21 SAR11 and 5 *Prochlorococcus* single-cell amplified genomes (SAGs). These SAGs represent reference genomes in an ocean region with sparse coverage: only one cultured *Prochlorococcus* genome (27) and two cultured SAR11 genomes (46) are currently available from the Red Sea. The SAR11 SAGs also represent genomes from clades without other sequenced representatives: two SAGs from subclade Ib and three SAGs from subclade IId (Fig. 1).

**Figure 1.**
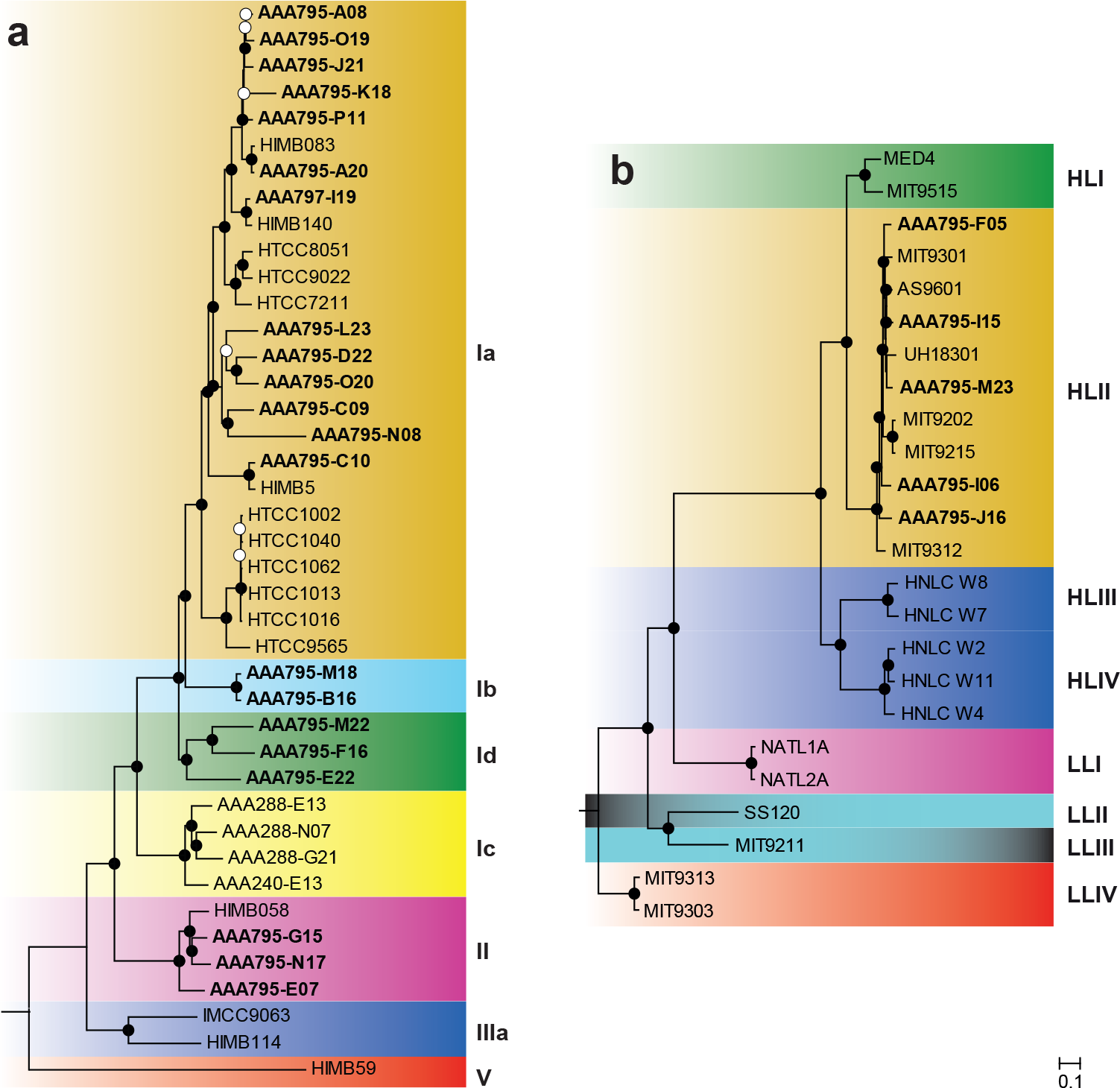
Maximum-likelihood proteomic trees for single-cell genomes from this study (bold), plus a representative set of cultured genomes. Trees were built from concatenated alignments of (a) 89 SAR11 and (b) 96 *Prochlorococcus* single-copy orthologous genes. Bootstrap values are indicated at the nodes (solid circles ≥80% and open circles ≥50%). Scale bars are equal to 0.1 changes per site. The Red Sea SAR11 SAGs cluster with subclades Ia, Ib, Id, and II. The Red Sea *Prochlorococcus* SAGs all cluster with ecotype HLII.

To account for and remove any possible contaminating DNA sequences, assembled contigs were retained only if they were part of a SAR11 or *Prochlorococcus* Metawatt bin or if they had a top-10 BLASTN hit to a *Prochlorococcus* or SAR11 genome (methods). In Metawatt, assignment to bins is based on tetranucleotide frequency, and the average taxonomy of the bin is determined by BLAST of 500-bp fragments of all the contigs against a prokaryotic database (35). A contig matching the tetranucleotide frequency of a SAR11 or *Prochlorococcus* bin could be retained even if it contained contradictory or missing taxonomic information information. However, to check if our secondary, BLASTN-based assignment process could be biased against short contigs, which might lack a neighboring anchor gene, we analyzed the distribution of contig lengths between retained and removed contigs for each SAG. We found that in most cases (20 of 26 SAGs) the median sizes of retained and removed contigs were not different (Fig. S2); in 6 SAGs the retained contigs were larger than the removed contigs (Mann–Whitney U, *p* < 0.05, two-tailed).

Genome size and completeness was greater for *Prochlorococcus* SAGs than SAR11 SAGs. Size of *Prochlorococcus* SAGs ranged from from 1.28–1.46 Mbp in 85–221 contigs, containing 1428–1710 genes; SAR11 SAGs ranged from 0.29–1.14 Mbp in 55–157 contigs, containing 342–1199 genes (Table 1). Completeness was calculated by two methods: fraction of single-copy core genes observed and CheckM completeness score; genome redundancy was calculated by CheckM. Completeness of *Prochlorococcus* SAGs ranged from 85.9–90.3% core completeness and 70.7–78.7% CheckM completeness; SAR11 SAGs ranged from 20.3–90.0% core completeness and 19.1–76.7% CheckM completeness (Table 1). Genome redundancy of *Prochlorococcus* SAGs ranged from 0.1–1.0%, and of SAR11 SAGs ranged from 0.0–1.4% (Table 1). Plotting the number of single-copy core genes as a function of total contig size (Fig. S3) showed a strong correlation between total contig size and number of single-copy core genes; this analysis illustrates the greater completeness of the *Prochlorococcus* SAGs relative to the SAR11 SAGs.

**Table 1.** Genomic features of *Prochlorococcus* and SAR11 single-cell genomes. Single cells were isolated from a surface sample from the Eastern Red Sea (19.75 °N, 40.05 °E). *Prochlorococcus* clades are ecotypes; SAR11 clades are subclades. Completeness is reported as the fraction of 1144 *Prochlorococcus* or 649 SAR11 single-copy core OGs found in each SAG; completeness is also reported as the percent of bacterial single-copy core OGs present as determined by CheckM. Redundancy of bacterial single-copy core OGs is defined as the “contamination” parameter from the CheckM software.

| Genus | SAG ref. no. | Clade | Contigs | Assembled size (bp) | Genes | Single-copy core genes | Completeness (core, %) | Completeness (CheckM, %) | Redundancy (CheckM, %) | G+C (%) |
| --- | --- | --- | --- | --- | --- | --- | --- | --- | --- | --- |
| *Prochlorococcus* | SCGC AAA795-F05 | HLII | 136 | 1,418,374 | 1632 | 1033 | 90.2 | 78.6 | 0.27 | 31.4 |
| *Prochlorococcus* | SCGC AAA795-I06 | HLII | 120 | 1,388,767 | 1604 | 981 | 85.9 | 77.5 | 0.10 | 31.1 |
| *Prochlorococcus* | SCGC AAA795-I15 | HLII | 221 | 1,282,941 | 1428 | 989 | 86.6 | 70.7 | 0.97 | 31.3 |
| *Prochlorococcus* | SCGC AAA795-J16 | HLII | 85 | 1,463,721 | 1691 | 1033 | 90.3 | 78.7 | 0.52 | 31.0 |
| *Prochlorococcus* | SCGC AAA795-M23 | HLII | 93 | 1,443,989 | 1710 | 1012 | 88.7 | 74.6 | 0.34 | 31.2 |
| SAR11 | SCGC AAA795-A08 | Ia | 61 | 374,567 | 384 | 158 | 24.3 | 24.5 | 0.00 | 28.3 |
| SAR11 | SCGC AAA795-A20 | Ia | 63 | 1,140,609 | 1199 | 584 | 90.0 | 76.7 | 0.00 | 29.1 |
| SAR11 | SCGC AAA795-B16 | Ib | 95 | 551,717 | 600 | 331 | 51.0 | 34.7 | 0.06 | 29.4 |
| SAR11 | SCGC AAA795-C09 | Ia | 82 | 667,038 | 734 | 390 | 60.1 | 44.6 | 0.88 | 28.4 |
| SAR11 | SCGC AAA795-C10 | Ia | 55 | 477,445 | 503 | 213 | 32.8 | 34.9 | 0.23 | 29.3 |
| SAR11 | SCGC AAA795-D22 | Ia | 68 | 1,010,421 | 1082 | 555 | 85.5 | 69.9 | 0.60 | 28.8 |
| SAR11 | SCGC AAA795-E07 | II | 101 | 681,366 | 737 | 418 | 64.4 | 56.9 | 1.37 | 29.7 |
| SAR11 | SCGC AAA795-E22 | Ib | 63 | 801,227 | 820 | 417 | 64.3 | 47.6 | 0.34 | 29.0 |
| SAR11 | SCGC AAA795-F16 | Ib | 74 | 945,491 | 1017 | 509 | 78.4 | 65.9 | 0.00 | 29.1 |
| SAR11 | SCGC AAA795-G15 | II | 62 | 294,337 | 342 | 132 | 20.3 | 19.1 | 0.46 | 30.5 |
| SAR11 | SCGC AAA795-J21 | Ia | 77 | 872,902 | 954 | 404 | 62.2 | 51.5 | 0.70 | 29.1 |
| SAR11 | SCGC AAA795-K18 | Ia | 114 | 731,292 | 782 | 314 | 48.4 | 48.7 | 0.70 | 29.9 |
| SAR11 | SCGC AAA795-L23 | Ia | 150 | 834,822 | 910 | 489 | 75.3 | 54.4 | 0.60 | 27.8 |
| SAR11 | SCGC AAA795-M18 | Ib | 61 | 1,050,527 | 1072 | 456 | 70.3 | 58.9 | 1.41 | 29.2 |
| SAR11 | SCGC AAA795-M22 | Ib | 80 | 860,157 | 921 | 515 | 79.4 | 64.2 | 0.13 | 29.4 |
| SAR11 | SCGC AAA795-N08 | Ia | 157 | 575,315 | 622 | 272 | 41.9 | 33.3 | 0.55 | 29.1 |
| SAR11 | SCGC AAA795-N17 | II | 94 | 611,592 | 620 | 361 | 55.6 | 38.0 | 0.42 | 29.5 |
| SAR11 | SCGC AAA795-O19 | Ia | 62 | 804,609 | 862 | 379 | 58.4 | 54.2 | 0.04 | 29.1 |
| SAR11 | SCGC AAA795-O20 | Ia | 62 | 1,009,143 | 1074 | 526 | 81.0 | 69.0 | 0.04 | 29.0 |
| SAR11 | SCGC AAA795-P11 | Ia | 127 | 977,727 | 1021 | 485 | 74.7 | 52.4 | 1.32 | 29.2 |
| SAR11 | SCGC AAA797-I19 | Ia | 77 | 1,016,895 | 1071 | 468 | 72.1 | 66.4 | 0.59 | 29.2 |

Taxonomic assignment of SAGs to clades was done by comparing SAGs against reference genomes using several methods. Phylogenetic analysis was done on concatenated proteins (89 SAR11 and 96 *Prochlorococcus* shared single-copy orthologous genes) using the maximum likelihood method (methods). Nucleotide composition (G+C content and *k*-mer composition) was calculated and compared to reference genomes. Ordination using principal components analysis (PCA) of *k*-mer composition and OG composition (presence–absence of each OG in each genome) was used to visualize SAGs in relation to known clades of SAR11 and *Prochlorococcus*.

Phylogenetic analysis of concatenated proteins (Fig. 1) revealed that *Prochlorococcus* SAGs were all ecotype HLII (5/5). Surveys of the Red Sea using 16S–23S rRNA internal transcribed spacer (ITS) amplicon sequencing (47), *rpoC1* gene amplicon sequencing (48), and metagenomic sequencing (12) have each shown that HLII is the dominant *Prochlorococcus* ecotype in the surface Red Sea. This pattern is consistent with temperature-driven ecotype distribution patterns of *Prochlorococcus*, where ecotype HLII is predominant in warm/tropical surface waters (and has a higher thermal tolerance in culture) and ecotype HLI is predominant in cool/subtropical surface waters (49). SAR11 SAGs were predominantly subclade Ia (13/21), with the remainder subclades Ib (2/21), Id (3/21), and II (3/21). Placement of the SAR11 SAGs in these respective clades is supported by a previous phylogenetic analysis of 16S rRNA gene sequences that included these SAGs (10). Surveys using amplicon sequencing of the 16S rRNA gene (50) and metagenomic sequencing (12) have both shown that SAR11 subclade Ia dominates the surface Red Sea. Subclade distributions in the 16S survey (50) approximately matched the distribution of the SAG subclades here, suggesting that the SAGs may approximate the natural SAR11 population.

DNA G+C content of the *Prochlorococcus* SAGs ranged from 31.0-31.4% (Table 1), which is typical of genomes of *Prochlorococcus* ecotype HLII (51). G+C content of the SAR11 SAGs was lower, ranging from 27.8-30.5% (Table 1). We have previously shown, using the SAR11 SAGs and other SAR11 genomes, that the ratio of nonsynonymous to synonymous nucleotide mutations and other genomic evidence in SAR11 genomes is consistent with selection for low nitrogen driving the low G+C content in marine SAR11 (10).

Ordination by PCA of genome properties provided visualization and in some cases improved resolution of genome taxonomy relative to tree-based methods. For nucleotide composition analysis, six-nucleotide words (6-mers) were chosen to balance computational time and information content. The distribution of all 4096 possible 6-mers across the genomes was subject to dimensionality reduction using PCA and plotted as the first two principal components (PCs). The first PC explains 27% and 67% of the variation, respectively, for the SAR11 genomes (Fig. 2a) and the *Prochlorococcus* genomes (Fig. 2b). The PCA plots show wider spread in the SAR11 genomes than in the *Prochlorococcus* genomes; both cluster by clade, but the *Prochlorococcus* genomes are more tightly clustered, with three main clusters (Fig. 2b): HLI nested within HLII and near HLIII/IV (lower-left), then LLI (middle-left) next-closest followed by LLII and LLIII (upper-left), and then LLIV distant from the others and more disperse (lower-right).

**Figure 2.**
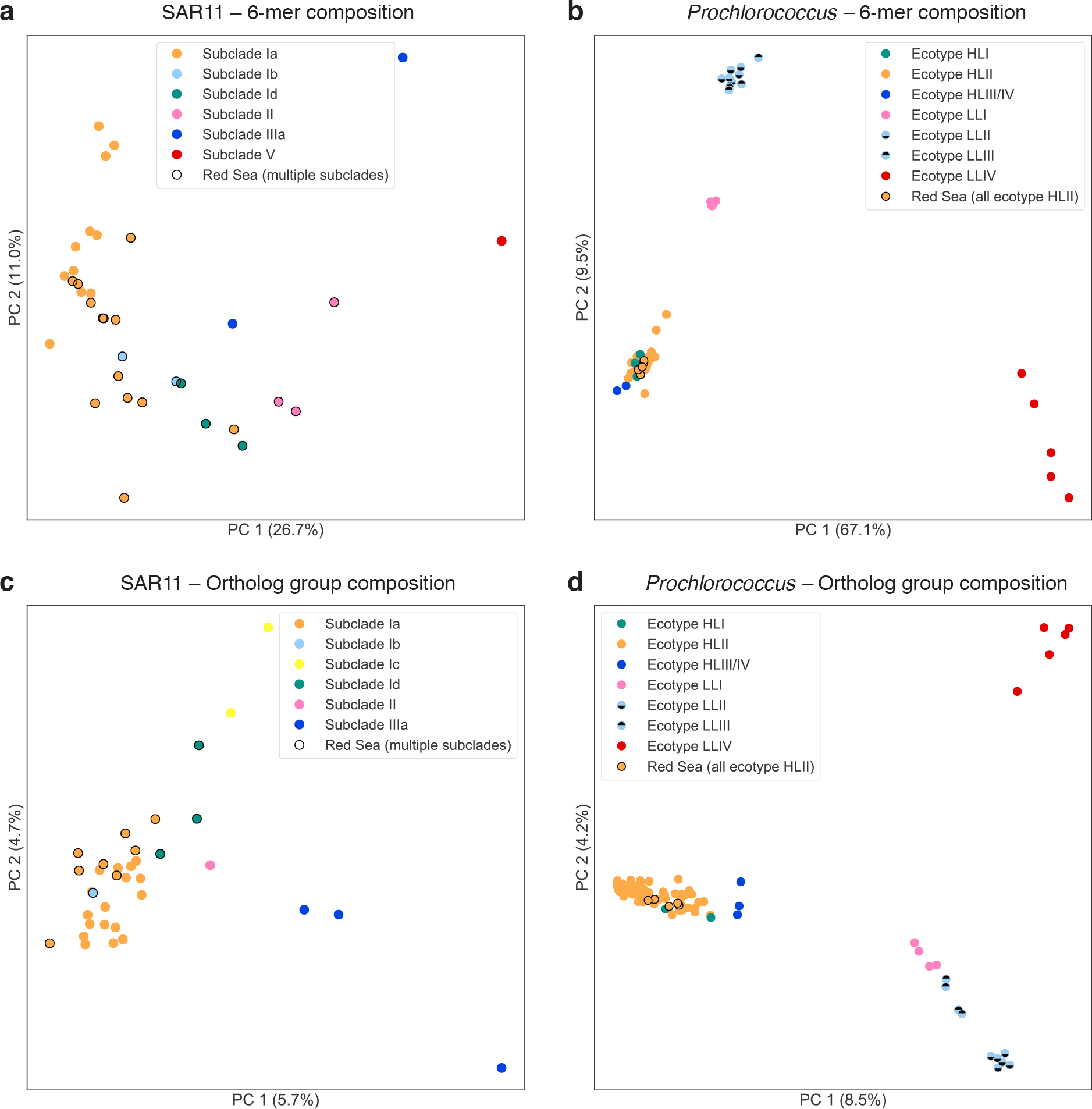
PCA ordination of SAGs and genomes based on (a, b) hexanucleotide (6-mer) composition and (c, d) ortholog group (OG) composition. Genomes are colored by clade; single-cell genomes from the Red Sea (this study) are circled in black. OG counts, prior to PCA ordination, were subsampled to 800 (SAR11) or 1400 (*Prochlorococcus*). While both nucleotide composition and OG composition cluster genomes into discrete groups by clade, OG composition differentiate clades more clearly, as exemplified by the separation of *Prochlorococcus* clades LLII and LLIII (panel d).

Ordination by PCA of OG composition was done following subsampling of OG counts down to 800 gene counts per SAR11 genome and 1400 gene counts per *Prochlorococcus* genome (methods). This had the effect of dropping 9 SAR11 SAGs, but it allowed the genomes to have even depth of coverage for PCA calculation. PCA ordination revealed patterns of OG composition of SAR11 genomes (Fig. 2c) and *Prochlorococcus* genomes (Fig. 2d). PC1 and PC2 each explained 6-9% of the variation for both sets of genomes. For SAR11, ordination of OG composition clustered by clade approximately as well as 6-mer composition. For *Prochlorococcus*, PCA of OG composition provided good separation of the low-light ecotypes (LLI, LLII, LLIII, and LLIV), whereas the high-light ecotypes HLI and HLII formed a single cluster with HLIII/IV nearby.

Of particular interest to investigations of the low-light adapted *Prochlorococcus* ecotypes, we note that OG composition clearly distinguished between genomes of ecotypes LLII and LLIII. It has previously been observed that phylogenetic analysis (ITS region) (52, 53) does not resolve ecotypes LLII and LLIII (identified as high B/A II and III by (54)). Similarly, our analysis of 6-mer composition also could not resolve these two low-light ecotypes. Our method of “OG ordination”, however, did distinguish these ecotypes. Thus it is helpful that OG distributions can assign genomes to ecotypes that are indistinguishable by other taxonomic or phylogenetic methods. The rich genotypic information provided by OG distribution patterns, combined with an ordination method like PCA, could be applied to other microbial groups for taxonomic classification of closely related genomes.

### Gene clustering and identification of Red-Sea-associated ortholog groups

The SAGs described here come from an undersampled region of the ocean (the Red Sea) and in part from undersampled clades of marine bacteria (SAR11 subclades Ib, Id, and II), and therefore provide the opportunity to identify OGs specific for these clades or possibly endemic to this ocean region. To investigate these patterns, we combined the Red Sea SAGs with available cultured genomes and SAGs (separately for *Prochlorococcus* and SAR11), clustered genes into OGs using a Markov clustering algorithm (OrthoMCL, methods), and identified those OGs found only in the Red Sea SAGs and/or only in certain clades.

We identified 878 SAR11 OGs and 96 *Prochlorococcus* Red-Sea-associated OGs (RS-OGs), that is, OGs not found (in this analysis) in genomes from other parts of the ocean (supplemental file 1). These totals represent 16.7% of all (19.1% of non-core) SAR11 OGs and 0.9% of all (1.0% of non-core) *Prochlorococcus* OGs. Many of the RS-OGs were found only in a single clade: 96 in Prochlorococcus ecotype HLII, 484 in SAR11 subclade Ia, 101 in SAR11 subclade Ib, 101 in SAR11 subclade Id, and 132 in SAR11 subclade II. The numerous clade-specific OGs present targets for understanding ecotype-specific physiology.

The first pattern of note was that there were more RS-OGs in the SAR11 SAGs than in the *Prochlorococcus* SAGs. This reflects the large contribution of our SAR11 SAGs to the sequenced SAR11 pangenome: the number of SAR11 Red Sea SAGs (=21) was nearly as many as the number of SAR11 reference genomes (=26). In contrast, the number of *Prochlorococcus* Red Sea SAGs (=5) was only 3% of the number of *Prochlorococcus* reference genomes (=140). Emphasizing the effect of the genome reference database on estimates of OG endemicity, after new *Prochlorococcus* genomes (9, 52) were added to the clustering, the number of RS-OGs dropped from 1192 to 96 (Fig. S4). Another explanation for the greater number of new SAR11 OGs is that the SAR11 SAGs span previously unsampled or undersampled clades: 334 of the 878 Red-Sea-associated SAR11 OGs were found in only one of subclade Ib, Id, or II. Furthermore, SAR11 is a broader phylogenetic group, based on 16S rRNA diversity, than *Prochlorococcus* (55), and therefore its pangenome may be expected to be larger. In summary, we suspect that the large number of new SAR11 OGs (=878), in general, more likely reflects the current dearth of sequence data for SAR11 rather than a significant degree endemism due to isolation and/or selection.

The second pattern we examined was inspired by our question about possible endemic gene content in the Red Sea: based on the geographic isolation of the Red Sea and its unique combination of physicochemical conditions (simultaneously high irradiance, high salinity, high temperature, and low nutrients), do genomes isolated from the Red Sea exhibit endemic OG content encoding adaptive functions for this environment? The answer that emerged to this question is that there were some indications of possible endemic adaptations to the Red Sea; however, there were no new pathways identifiable, most of the OGs with annotated functions were found in only one or two SAGs, and the majority of OGs encoded hypothetical proteins with no assigned function.

The majority of RS-OGs were hypothetical proteins: 82% (723 of 878) for SAR11 and 91% (87 of 96) for *Prochlorococcus*. It was difficult to infer possible adaptive functions for OGs with no predicted functions; however, these OGs may be referenced later when new approaches for annotating conserved hypotheticals are developed. The remaining non-hypothetical OGs (155 SAR11, 9 *Prochlorococcus*), i.e., those with predicted functions, are listed in Table S2. While we could not detect a widespread signature of adaptation to the Red Sea environment—i.e., RS-OGs with annotated functions represented across multiple SAGs—below we highlight a few sparsely represented RS-OGs that may have adaptive functionality in the Red Sea environment, some with possible biotechnological potential.

Among *Prochlorococcus* SAGs, none of the 9 non-hypothetical RS-OGs (Table S2) were found in more than one SAG. One OG (proch20425) found in SCGC AAA795-M23 encodes UvrABC system protein B, responsible for repair of DNA damage. We could posit that this enzyme is found preferentially in the Red Sea because of the year-round high irradiance, which increases the rate of DNA damage in cells.

Among SAR11 SAGs, there were 21 non-hypothetical RS-OGs found in two or more SAGs and another 134 found in only one SAG (Table S2). These OGs show links to high light adaptation, motility, and nitrogen and phosphorus assimilation. One OG (pelag14710, found in one SAG) encodes a photolyase enzyme that repairs damaged DNA caused by exposure to ultraviolet light. Pyrophosphatase (pelag15064, found in one SAG) is involved in the hydrolysis of inorganic pyrophosphate into two orthophosphates and may have a role in phosphorus utilization. Allantoinase (pelag15247) and urease accessory protein UreF (pelag14490) are each found in one SAR11 SAG. These enzymes involved in phosphorus and nitrogen metabolism may provide an adaptive advantage in the Red Sea, which exhibits co-limitation to both elements and may be relatively more nitrogen-limited (12, 15). Several of the SAR11 RS-OGs encode enzymes with biotechnological relevance. DNA polymerase I (pelag12679, pelag14776, pelag14807) from this higher temperature environment could have heat-resistant properties, for example, marginal thermostability conferred by amino acid substitutions (56).

After the major analyses had been completed for this study, two SAR11 genomes (46) and one *Prochlorococcus* genome (27) derived from cultivated strains were sequenced, and four *Prochlorococcus* genomes were assembled from metagenomes (57). Of the SAR11 genomes, one was assigned to subclade Ia and the other to subclade Ib (46). Of note, the subclade Ia genome (RS39) contained several OGs also found among the Red-Sea-associated SAR11 OGs: 3-oxoacyl-acyl-carrier-protein synthase, ABC branched amino acid transporter, arylsulfotransferase, formate dehydrogenases, glycosyl transferases, methyltransferases, sialic acid synthase, sucrose synthase, sulfotransferases, and a type II restriction–modification system. Several of these functions may play roles in one-carbon and sugar metabolism by SAR11 in the Red Sea (46). The *Prochlorococcus* genome was assigned to the HLII ecotype and notably contained a pathway for biosynthesis of the osmolyte (compatible solute) glucosylglycerol (27). This pathway represents a possible adaptation to the higher salinity of the Red Sea. However, the three genes in this pathway were not found among the Red-Sea-associated *Prochlorococcus* OGs, nor were they found elesewhere among the retained or removed contigs from the Red Sea SAGs (BLASTN).

### Expression of ortholog groups in the Red Sea water column

To further test the idea that there could be OGs of ecological importance endemic to the Red Sea, we analyzed metatranscriptomes from the Red Sea. Any OGs with functional roles would be expected to be expressed in the Red Sea water column. We collected seawater and filtered the prokaryotic fraction from a station in the central Red Sea at four depths and 13 timepoints over a 48-hour period. We extracted and sequenced RNA from these samples, and mapped the reads to the Red Sea SAGs.

We found that a majority of RS-OGs were expressed in one or more sample (64% SAR11, 66% *Prochlorococcus*; Fig. 3a,b). This was more than the fraction of non-RS-OGs expressed (32% SAR11, 20% *Prochlorococcus*; Fig. 3c,d). We were curious if the high fraction of non-RS-OGs that were unexpressed was due to many of these OG being singletons (OGs having only one member). To the contrary, heatmaps of OG size vs. number of metatranscriptomes in which the OG was found (Fig. 3, inset) do not show a high density of singleton OGs having no expression in non-RS-OGs, and rather the trend toward singletons is more common in RS-OGs.

**Figure 3.**
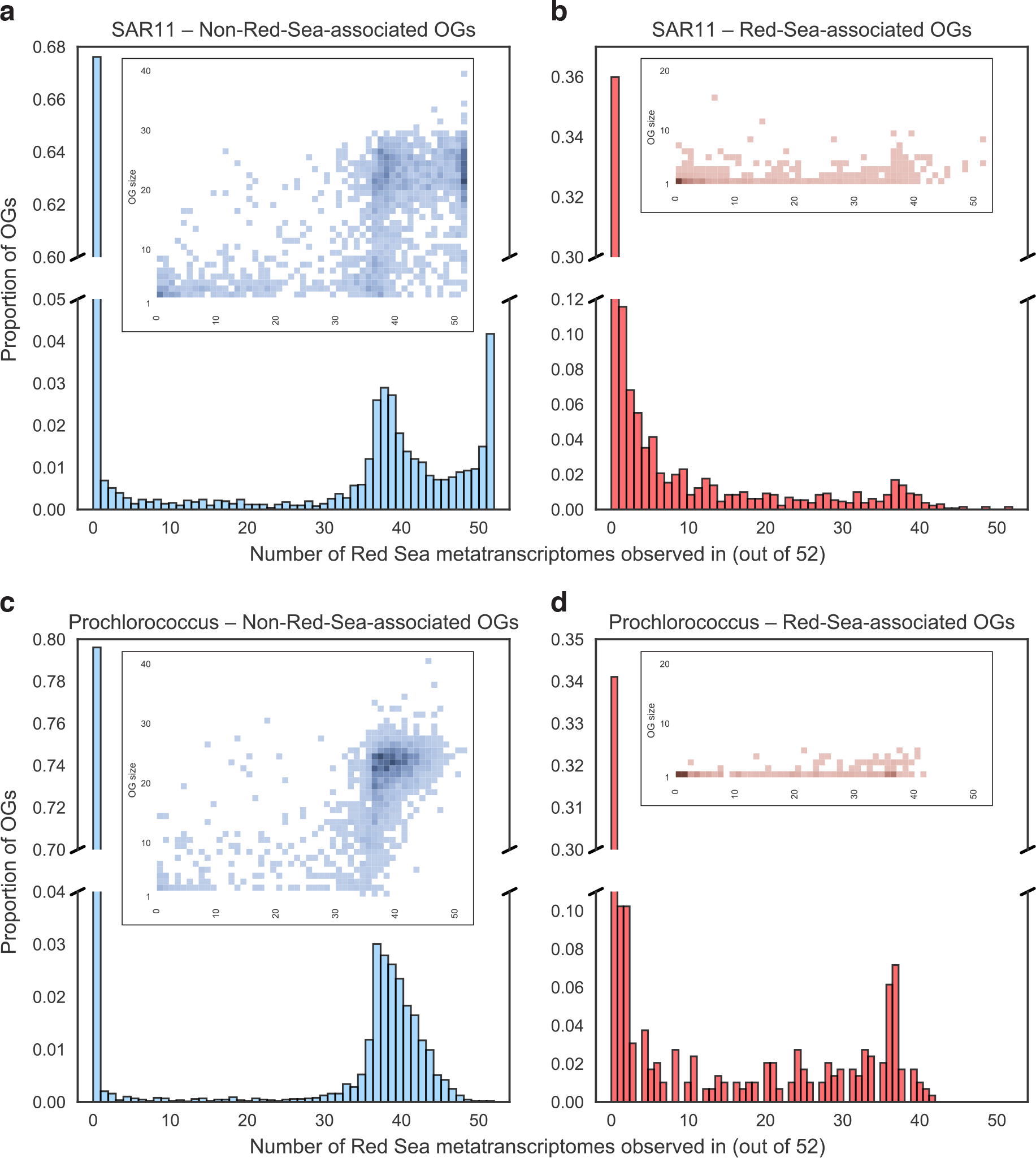
Expression of SAG ortholog groups (OGs) in Red Sea metatranscriptomes. The 52 metatranscriptomes span four depths and 13 timepoints over a 48-hour period (every 4 hours) from a station in the central Red Sea. Histograms show the number of metatranscriptomes found in of (a) SAR11 non-RS-OGs, (b) SAR11 RS-OGs, (c) *Prochlorococcus* non-RS-OGs, and (d) *Prochlorococcus* RS-OGs. Heatmaps (inset) show the density of OGs based on OG size (number of total copies across the SAGs) and the number of metatranscriptomes an OG is found in. RS-OGs were more likely than other OGs to be expressed in one or more sample, and non-RS-OGs that were expressed were more likely to be expressed in a high number of samples.

Of OGs expressed in at least one sample, non-RS-OGs (Fig. 3a,c) tended to be expressed in more samples than RS-OGs (Fig. 3b,d). This is consistent with many of the non-RS-OGs being core genes, many of which are housekeeping genes that are often constitutively expressed. Overall, the expression patterns indicate that the majority of RS-OGs are transcribed to messenger RNA, consistent with the synthesis of functional gene products.

### Distribution of ortholog groups across the global ocean

The analysis to this point has focused on the distribution of OGs among cultured and single-cell genomes and their expression in the Red Sea water column. A set of OGs has been found that is exclusive to Red Sea genomes (to date), and a majority of them are expressed in the water column. However, we cannot rule out the possibility that these OGs appear endemic only because more genomes are not available from around the World Ocean. If we extended our search to global marine metagenomes, instead of just genomes, would we in fact find these putative endemic OGs in other seas?

To investigate the possibility that, contrary to our original hypothesis, there may be few truly endemic OGs in the Red Sea microbial community, we analyzed metagenomes collected from across the global ocean by the *Tara* Oceans expedition. We searched for SAR11 and *Prochlorococcus* OGs in 139 prokaryote-fraction metagenomes from the *Tara* Oceans expedition (44), which come from several depths in the water column: surface, mixed layer, deep chlorophyll maximum, and mesopelagic zone. We queried the dataset to determine what fraction of all OGs and what fraction of RS-OGs could be found outside the Red Sea. If RS-OGs represent endemic gene content of the Red Sea, we would expect to find them absent from metagenomes from other regions. Our approach was complementary to a recent study that analyzed the global metapangenome of *Prochloroccocus* in the *Tara* metagenomes, showing the distributions of gene clusters (OGs) with strain-level resolution across the *Tara* samples (58). In the work here, we employed rarefaction and ordination techniques, with a particular focus on RS-OGs.

The presence or absence of SAR11 and *Prochlorococcus* orthologs in *Tara* Oceans prokaryote-fraction metagenomes (supplemental files 7 and 8) was plotted as rarefaction curves (Fig. 4). *Tara* Oceans metagenomes were added randomly one by one, and the fraction of SAR11 and *Prochlorococcus* OGs found was tallied and plotted. The rarefaction curves show the average ± standard deviation of 1000 permutations. They also show the best-case (and worst-case) scenarios, that is, the fraction of OGs found if each new metagenome adds the most (or fewest) new OGs. Between 70–85% of OGs could be found in one or more *Tara* Oceans metagenome (Fig. 4), and in the best-case scenarios it took at most ten metagenomes to find 90% of these OGs (Table S3). The percentage of OGs not found (15–30%) was independent of whether they were ‘Red-Sea-associated’ or not. This result combined with the rarefaction analysis suggests these OGs would be unlikely to be found in the *Tara* samples with deeper sequencing. It is possible that some OGs may be rare and/or divergent enough to be undetectable with the current methodological approach.

**Figure 4.**
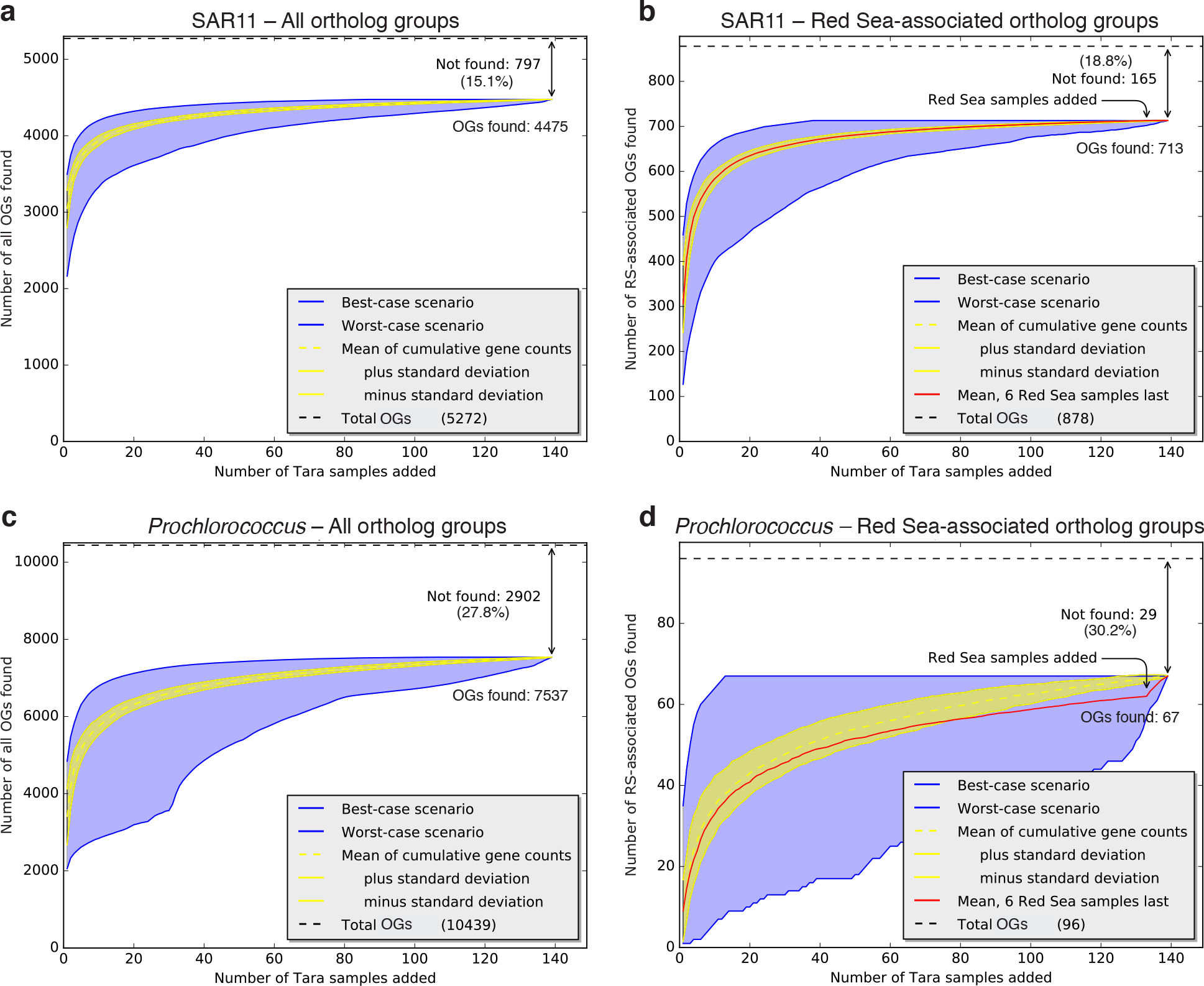
Rarefaction analysis showing the proportion of (a, c) all OGs and (b, d) RS-OGs of SAR11 and *Prochlorococcus* observed in *Tara* Oceans metagenome samples. Curves show the cumulative number of OGs observed in *Tara* Oceans samples (e-value < 1e-5) as more samples are added. Yellow lines show the average ± standard deviation of 1000 permutations of randomly added samples. Blue lines show the “best-case scenario” (each sample added is that with the most number of new OGs observed) and “worst-case scenario” (each sample added is that with the fewest number of new OGs observed). Red lines show the mean of 1000 permutations of randomly added samples but with Red Sea samples (031_SRF_0.22-1.6, 032_DCM_0.22-1.6, 032_SRF_0.22-1.6, 033_SRF_0.22-1.6, 034_DCM_0.22-1.6, 034_SRF_0.22-1.6) added last. As more *Tara* metagenome samples are added to the analysis, the number of new OGs identified approaches a plateau where new samples do not reveal many new OGs. The same is true with RS-OGs, even when samples from the Red Sea are added last, with the exception of 5 *Prochlorococcus* OGs (proch20367, proch20368, proch20390, proch20423, and proch20438).

Across the 139 *Tara* Oceans prokaryote-fraction metagenomes, we found 84.9% (4475/5272) of all SAR11 OGs in one or more metagenome (leaving 15.1% not found; Fig. 4a) and 72.2% (7537/10439) of all *Prochlorococcus* OGs in one or more metagenome (leaving 27.8% not found; Fig. 4c). In the best-case scenarios, it took only 5 metagenomes to find 90% of the ‘found’ SAR11 OGs and 50 metagenomes to find 99%; it took only 10 metagenomes to find 90% of the ‘found’ *Prochlorococcus* OGs and 60 metagenomes to find 99% (Table S3). The fractions of OGs found were similar for RS-OGs, where 81.2% (713/878) of SAR11 OGs were found (leaving 18.8% not found; Fig. 4b) and 69.8% (67/96) of *Prochlorococcus* OGs were found (leaving 30.2% not found; Fig. 4d). That is, RS-OGs were about as likely to be found across the World Ocean as non-RS-OGs. For both SAR11 (Fig. S5a) and *Prochlorococcus* (Fig. S5b), considering the number of *Tara* metagenomes in which each OG was found, RS-OGs were less likely to be found in a large fraction of metagenomes, relative to all OGs. This is not surprising: the set of non-RS-OGs contains all of the core OGs, which would be expected to be found in most if not all samples.

To evaluate whether *Tara* Red Sea metagenomes contained any RS-OGs not already found in the non-Red Sea metagenomes, we tested scenarios where the Red Sea metagenomes were added last in the rarefaction analysis. There was no change in the mean curve of cumulative SAR11 OGs found when the six *Tara* Red Sea metagenomes were added last (Fig. 4b): all of the SAR11 RS-OGs could be found without examining the Red Sea metagenomes. In contrast, there were five *Prochlorococcus* RS-OGs that were added to the cumulative total when the *Tara* Red Sea metagenomes were added last (Fig. 4d). These five OGs, all with unknown function, represent a small fraction of the total Prochlorococcus pangenome (10439 OGs total). Given the available genomes, this study may have uncovered a small set of OGs (Table S2) that possibly reflect gene content endemic to or generally associated with Red Sea environmental conditions, and this marks an area for further research. In light of this metagenomic analysis, however, it appears that the putative RS-OGs provide a relatively minor contribution to the whole and that these new SAR11 and *Prochlorococcus* genomes from the Red Sea generally reflect global pangenomes.

Finally, we were curious if OG composition as a whole could show the Red Sea metagenomes to be different from the other metagenomes, despite the lack of evidence of endemic OGs. More generally, could the relative abundance of OGs across *Tara* be used to distinguish populations of *Prochlorococcus* and SAR11?

We used the tables of OG counts in the 63 *Tara* surface (SRF) prokaryote-fraction metagenomes to do PCA ordination on the *Tara* metagenomes (Fig. 5; top OGs driving separation among the metagenomes provided in Table S4). SAR11 OG composition (Fig. 5a) was not obviously structured by temperature differences in the temperate and tropical ranges, though Red Sea samples clustered together, and polar samples were separate from the others. *Prochlorococcus* OG composition (Fig. 5b), however, was structured by temperature differences in the temperate and tropical ranges. The four Red Sea samples were split, with two samples clustering with the warm samples and two samples with the cooler samples. These Red Sea samples are positioned where they would be expected based on temperature: the two southern samples (latitude: 18.4 °N, 22.0 °N) were warmer (temperature: 27.6 °C, 27.3 °C) and clustered with other warm/tropical samples (left side of PC1 in Fig. 5b); the two northern samples (latitude: 23.36 °N, 27.16 °N) were cooler (temperature: 25.8 °C, 25.1 °C) and clustered closer to the cool/temperate samples (right side of PC1 in Fig. 5b). Note these temperatures are lower than average Red Sea surface waters because the *Tara* Red Sea samples were collected in winter (January); by contrast, the Red Sea samples in the World Ocean Database (see above) were collected in spring (April). Given that temperature tolerances generally lack known genetic markers (59), these data suggest an area for future investigation.

**Figure 5.**
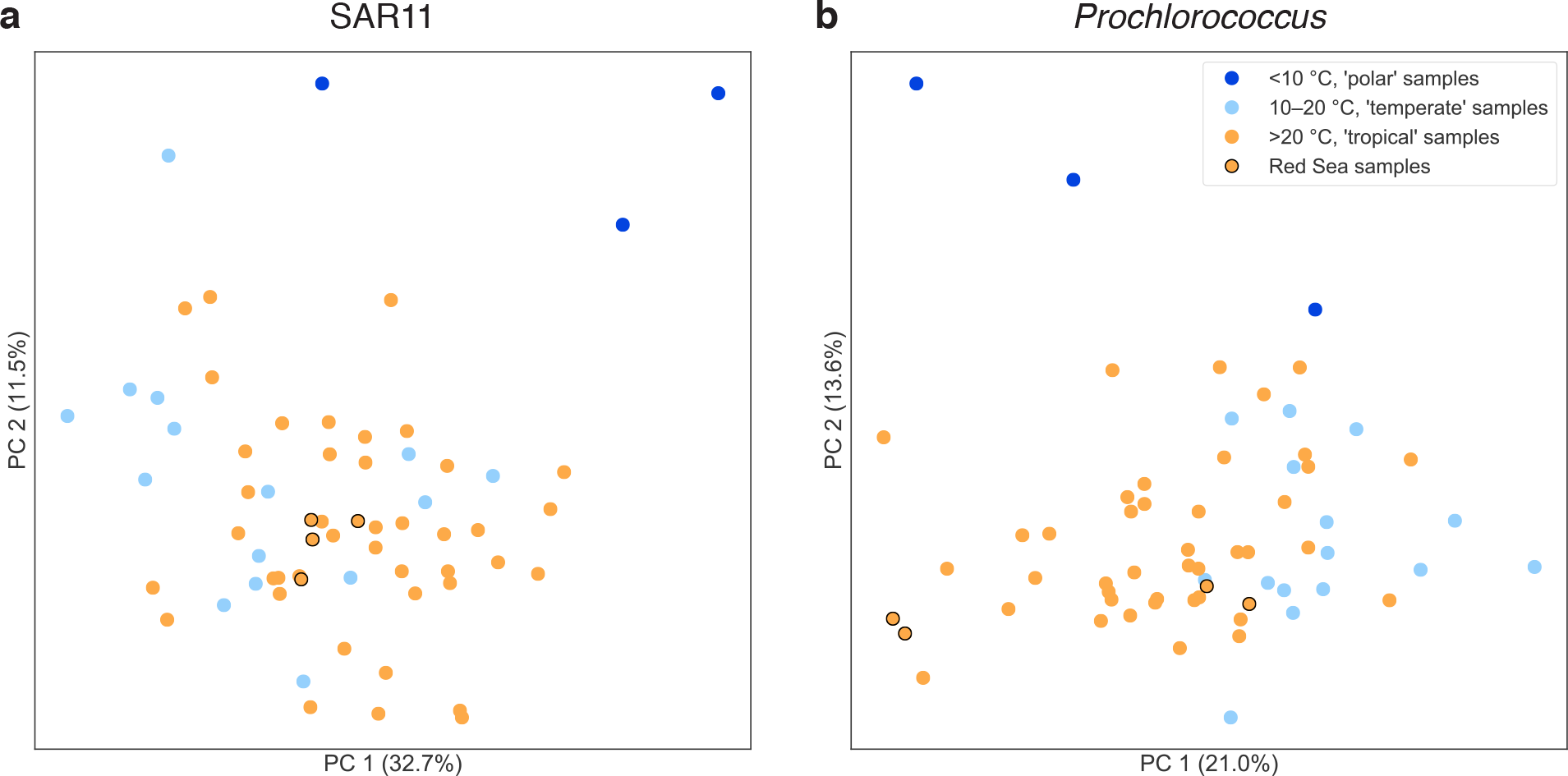
Principal components analysis of *Tara* Oceans surface samples by the abundance of (a) SAR11 and (b) *Prochlorococcus* OGs. The ordination shows the similarity of *Tara* Oceans samples to each other along the first two principal components. Samples are colored by *Tara* temperature categories: ‘polar’ samples (<10 °C) are dark blue, ‘temperate’ samples (10-20 °C) are light blue, ‘tropical’ samples (>20 °C) are orange, and Red Sea ‘tropical’ samples are orange with black edges. Red Sea samples and *Tara* samples generally show more separation based on temperature when ordinated by *Prochlorococcus* OG composition than by SAR11 OG composition.

In summary, the analysis of *Prochlorococcus* and SAR11 OGs in *Tara* Oceans metagenomes shows that (i) most “Red-Sea-associated” OGs are actually widely distributed across the World Ocean, not endemic to the Red Sea; and (ii) OG distribution patterns as a whole, taking relative abundance into account, place the Red Sea on a continuum with other seas, with patterns explained by environmental factors including temperature. Supporting this idea, differences in the relative abundance of OGs—with physicochemical properties covarying with OG functions—have been observed among the North Pacific, Sargasso Sea, Mediterranean Sea, and Red Sea in previous comparative metagenomics studies (11, 12). Despite the Red Sea existing at the periphery of multiple physicochemical parameters in the World Ocean, its distinctiveness may best be revealed by the relative abundance of OGs rather than in the wholesale presence or absence of OGs. In addition to this general pattern, this effort also identified a small set of putative and non-hypothetical proteins that warrant further ecological and biotechnological study.

### Conclusions and future directions

Here we analyzed SAR11 and *Prochlorococcus* SAGs from an undersampled ocean region, the Red Sea. This single-cell sequencing effort included SAR11 SAGs from undersampled clades and provided the first genomes from SAR11 subclades 1b and 1d. Our analysis of these genomes provided significant contributions to the reference databases of these organisms, adding 878 new ortholog groups to the SAR11 pangenome and 96 new ortholog groups to the *Prochlorococcus* pangenome. We described a new method called “OG ordination” that uses PCA of ortholog group composition to resolve phylogenetic differences in closely related genomes and used it to distinguish *Prochlorococcus* ecotypes LLII and LLIII in our samples.

How marine microbes are able to respond to a changing ocean will be critical to understanding the future biosphere of planet Earth. At the population and community levels, the cosmopolitan distribution of genetic functions may confer an advantage, enabling marine microbial populations and communities, as a whole, to rapidly respond and adapt to changing ocean conditions. Here we generally considered the Baas Becking hypothesis (“Everything is everywhere, but the environment selects”) from the perspective of gene ortholog groups (“Every OG is everywhere, but the environment selects”). The overall data analysis lends support to the Baas Becking hypothesis as applied to OGs. We described a small set of OGs that may be related to Red Sea environmental conditions and that mark areas for further investigation. However, the overall analysis was not consistent with endemism as a primary feature. Instead, we found Red Sea OGs to be nearly as prevalent across global ocean metagenomes as in Red Sea metagenomes. This view was supported by analysis of OG relative abundance rather than absolute presence-absence of OGs. Perhaps OGs may be present but undetectable in a region, and they become detectable after OG frequencies increase in response to environmental conditions (via the growth of cells containing those OGs). Therefore, genomic adaptations in a given ocean region may not simply reflect the presence of OGs unique to a region, but rather the relative abundance of generally cosmopolitan OGs.

## Acknowledgements

We thank Haiwei Luo for assistance building genome trees, Mamoon Rashid for consultation about decontamination methods, and Ramunas Stepanauskas and Nicole Poulton for assistance with the single-cell genomics protocol.

## References

1. Frias-Lopez J, Shi Y, Tyson GW, Coleman ML, Schuster SC, Chisholm SW, DeLong EF. 2008. Microbial community gene expression in ocean surface waters. Proc Natl Acad Sci USA 105:3805–3810.

2. Waldbauer JR, Rodrigue S, Coleman ML, Chisholm SW. 2012. Transcriptome and proteome dynamics of a light-dark synchronized bacterial cell cycle. PLoS ONE 7:e43432.

3. Coleman ML, Chisholm SW. 2007. Code and context: Prochlorococcus as a model for cross-scale biology. Trends Microbiol 15:398–407.

4. Good BH, McDonald MJ, Barrick JE, Lenski RE, Desai MM. 2017. The dynamics of molecular evolution over 60,000 generations. Nature 551:45–50.

5. Ochman H, Lawrence JG, Groisman EA. 2000. Lateral gene transfer and the nature of bacterial innovation. Nature 405:299–304.

6. Rocap G, Larimer FW, Lamerdin J, Malfatti S, Chain P, Ahlgren NA, Arellano A, Coleman M, Hauser L, Hess WR, Johnson ZI, Land M, Lindell D, Post AF, Regala W, Shah M, Shaw SL, Steglich C, Sullivan MB, Ting CS, Tolonen A, Webb EA, Zinser ER, Chisholm SW. 2003. Genome divergence in two Prochlorococcus ecotypes reflects oceanic niche differentiation. Nature 424:1042–1047.

7. Rodrigue S, Malmstrom RR, Berlin AM, Birren BW, Henn MR, Chisholm SW. 2009. Whole genome amplification and de novo assembly of single bacterial cells. PLoS ONE 4:e6864.

8. Grote J, Thrash JC, Huggett MJ, Landry ZC, Carini P, Giovannoni SJ, Rappé MS. 2012. Streamlining and core genome conservation among highly divergent members of the SAR11 clade. mBio 3:e00252–12.

9. Kashtan N, Roggensack SE, Rodrigue S, Thompson JW, Biller SJ, Coe A, Ding H, Marttinen P, Malmstrom RR, Stocker R, Follows MJ, Stepanauskas R, Chisholm SW. 2014. Single-cell genomics reveals hundreds of coexisting subpopulations in wild Prochlorococcus. Science 344:416–420.

10. Luo H, Thompson LR, Stingl U, Hughes AL. 2015. Selection Maintains Low Genomic GC Content in Marine SAR11 Lineages. Mol Biol Evol 32:2738–2748.

11. Coleman ML, Chisholm SW. 2010. Ecosystem-specific selection pressures revealed through comparative population genomics. Proc Natl Acad Sci USA 107:18634–18639.

12. Thompson LR, Field C, Romanuk T, Ngugi D, Siam R, El Dorry H, Stingl U. 2013. Patterns of ecological specialization among microbial populations in the Red Sea and diverse oligotrophic marine environments. Ecol Evol 3:1780–1797.

13. Berube PM, Biller SJ, Kent AG, Berta-Thompson JW, Roggensack SE, Roache-Johnson KH, Ackerman M, Moore LR, Meisel JD, Sher D, Thompson LR, Campbell L, Martiny AC, Chisholm SW. 2015. Physiology and evolution of nitrate acquisition in Prochlorococcus. ISME J 9:1195–1207.

14. Edwards FJ. 1987. Climate and oceanography, pp. 45–68. In Edwards, AJ, Head, SM (eds.), Key environments: Red sea. Pergamon, Oxford.

15. Post AF. 2005. Nutrient limitation of marine cyanobacteria, pp. 87–107. In Huisman, J, Matthijs, HCP, Visser, PM (eds.), Harmful cyanobacteria. Springer.

16. Thompson LR, Williams GJ, Haroon MF, Shibl A, Larsen P, Shorenstein J, Knight R, Stingl U. 2016. Metagenomic covariation along densely sampled environmental gradients in the Red Sea. ISME J 11:138–151.

17. Baas Becking LGM. 1934. Geobiologie of inleiding tot de milieukunde. W.P. Van Stockum & Zoon, The Hague, Netherlands.

18. Gibbons SM, Caporaso JG, Pirrung M, Field D, Knight R, Gilbert JA. 2013. Evidence for a persistent microbial seed bank throughout the global ocean. Proc Natl Acad Sci USA.

19. Gonnella G, Böhnke S, Indenbirken D, Garbe-Schönberg D, Seifert R, Mertens C, Kurtz S, Perner M. 2016. Endemic hydrothermal vent species identified in the open ocean seed bank. Nat Microbiol 1:16086.

20. Stepanauskas R, Sieracki ME. 2007. Matching phylogeny and metabolism in the uncultured marine bacteria, one cell at a time. Proc Natl Acad Sci USA 104:9052–9057.

21. Lane DJ, Pace B, Olsen GJ, Stahl DA, Sogin M, Pace NR. 1985. Rapid determination of 16S ribosomal RNA sequences for phylogenetic analyses. Proc Natl Acad Sci USA 82:6955–6959.

22. Page KA, Connon SA, Giovannoni SJ. 2004. Representative Freshwater Bacterioplankton Isolated from Crater Lake, Oregon. Appl Environ Microbiol 70:6542–6550.

23. Massana R, Murray AE, Preston CM, Delong EF. 1997. Vertical distribution and phylogenetic characterization of marine planktonic Archaea in the Santa Barbara Channel. Appl Environ Microbiol 63:50–56.

24. Béjà O, Suzuki MT, Heidelberg JF, Nelson WC, Preston CM, Hamada T, Eisen JA, Fraser CM, DeLong EF. 2002. Unsuspected diversity among marine aerobic anoxygenic phototrophs. Nature 415:630–633.

25. Stewart FJ, Dalsgaard T, Young CR, Thamdrup B, Revsbech NP, Ulloa O, Canfield DE, DeLong EF. 2012. Experimental incubations elicit profound changes in community transcription in OMZ bacterioplankton. PLoS ONE 7:e37118.

26. Markowitz VM, Mavromatis K, Ivanova NN, Chen I-MA, Chu K, Kyrpides NC. 2009. IMG ER: a system for microbial genome annotation expert review and curation. Bioinformatics (Oxford, England) 25:2271–2278.

27. Shibl AA, Ngugi DK, Talarmin A, Thompson LR, Blom J, Stingl U. 2018. The genome of a novel isolate of Prochlorococcus from the Red Sea contains transcribed genes for compatible solute biosynthesis. FEMS Microbiology Ecology.

28. Bolger AM, Lohse M, Usadel B. 2014. Trimmomatic: a flexible trimmer for Illumina sequence data. Bioinformatics (Oxford, England) 30:2114–2120.

29. Langmead B, Trapnell C, Pop M, Salzberg SL. 2009. Ultrafast and memory-efficient alignment of short DNA sequences to the human genome. CORD Conference Proceedings 10:R25–R25.

30. Nikolenko SI, Korobeynikov AI, Alekseyev MA. 2013. BayesHammer: Bayesian clustering for error correction in single-cell sequencing. BMC Genomics 14 Suppl 1:S7.

31. Bankevich A, Nurk S, Antipov D, Gurevich AA, Dvorkin M, Kulikov AS, Lesin VM, Nikolenko SI, Pham S, Prjibelski AD, Pyshkin AV, Sirotkin AV, Vyahhi N, Tesler G, Alekseyev MA, Pevzner PA. 2012. SPAdes: A New Genome Assembly Algorithm and Its Applications to Single-Cell Sequencing. J Comput Biol 19:455–477.

32. Kopylova E, Noé L, Touzet H. 2012. SortMeRNA: fast and accurate filtering of ribosomal RNAs in metatranscriptomic data. Bioinformatics (Oxford, England) 28:3211–3217.

33. Assefa S, Keane TM, Otto TD, Newbold C, Berriman M. 2009. ABACAS: algorithm-based automatic contiguation of assembled sequences. Bioinformatics (Oxford, England) 25:1968–1969.

34. Bonfield JK, Smith KF, Staden R. 1995. A new DNA sequence assembly program. Nucleic Acids Res 23:4992–4999.

35. Strous M, Kraft B, Bisdorf R. 2012. The binning of metagenomic contigs for microbial physiology of mixed cultures. Front Microbiol.

36. Aziz RK, Bartels D, Best AA, DeJongh M, Disz T, Edwards RA, Formsma K, Gerdes S, Glass EM, Kubal M, Meyer F, Olsen GJ, Olson R, Osterman AL, Overbeek RA, McNeil LK, Paarmann D, Paczian T, Parrello B, Pusch GD, Reich C, Stevens R, Vassieva O, Vonstein V, Wilke A, Zagnitko O. 2008. The RAST Server: rapid annotations using subsystems technology. BMC Genomics 9:75.

37. Li L, Stoeckert CJ, Roos DS. 2003. OrthoMCL: identification of ortholog groups for eukaryotic genomes. Genome Res 13:2178–2189.

38. Parks DH, Imelfort M, Skennerton CT, Hugenholtz P, Tyson GW. 2015. CheckM: assessing the quality of microbial genomes recovered from isolates, single cells, and metagenomes. Genome Res 25:1043–1055.

39. Contreras-Moreira B, Vinuesa P. 2013. GET_HOMOLOGUES, a versatile software package for scalable and robust microbial pangenome analysis. Appl Environ Microbiol 79:7696.

40. Katoh K. 2005. MAFFT version 5: improvement in accuracy of multiple sequence alignment. Nucleic Acids Res 33:511–518.

41. Lanfear R, Calcott B, Ho SYW, Guindon S. 2012. Partitionfinder: combined selection of partitioning schemes and substitution models for phylogenetic analyses. Mol Biol Evol 29:1695–1701.

42. Stamatakis A. 2014. RAxML version 8: a tool for phylogenetic analysis and post-analysis of large phylogenies. Bioinformatics (Oxford, England).

43. Pedregosa F, Varoquaux G, Gramfort A, Michel V, Thirion B, Grisel O, Blondel M, Prettenhofer P, Weiss R, Dubourg V, Vanderplas J, Passos A, Cournapeau D, Brucher M, Perrot M, Duchesnay É. 2011. Scikit-learn: machine learning in Python. J Mach Learn Res 12:2825–2830.

44. Sunagawa S, Coelho LP, Chaffron S, Kultima JR, Labadie K, Salazar G, Djahanschiri B, Zeller G, Mende DR, Alberti A, Cornejo-Castillo FM, Costea PI, Cruaud C, d’Ovidio F, Engelen S, Ferrera I, Gasol JM, Guidi L, Hildebrand F, Kokoszka F, Lepoivre C, Lima-Mendez G, Poulain J, Poulos BT, Royo-Llonch M, Sarmento H, Vieira-Silva S, Dimier C, Picheral M, Searson S, Kandels-Lewis S, Bowler C, Vargas C de, Gorsky G, Grimsley N, Hingamp P, Iudicone D, Jaillon O, Not F, Ogata H, Pesant S, Speich S, Stemmann L, Sullivan MB, Weissenbach J, Wincker P, Karsenti E, Raes J, Acinas SG, Bork P. 2015. Ocean plankton. Structure and function of the global ocean microbiome. Science 348:1261359–1261359.

45. Buchfink B, Xie C, Huson DH. 2014. Fast and sensitive protein alignment using DIAMOND. Nat Meth 12:59–60.

46. Jimenez-Infante F, Ngugi DK, Vinu M, Blom J, Alam I, Bajic VB, Stingl U. 2017. Genomic characterization of two novel SAR11 isolates from the Red Sea, including the first strain of the SAR11 Ib clade. FEMS Microbiol Ecol 93.

47. Shibl AA, Thompson LR, Ngugi DK, Stingl U. 2014. Distribution and diversity of Prochlorococcus ecotypes in the Red Sea. FEMS Microbiol Lett 356:118–126.

48. Shibl AA, Haroon MF, Ngugi DK, Thompson LR, Stingl U. 2016. Distribution of Prochlorococcus Ecotypes in the Red Sea Basin Based on Analyses of rpoC1 Sequences. Front Mar Sci 3.

49. Johnson ZI, Zinser ER, Coe A, McNulty NP, Woodward EMS, Chisholm SW. 2006. Niche partitioning among Prochlorococcus ecotypes along ocean-scale environmental gradients. Science 311:1737–1740.

50. Ngugi DK, Stingl U. 2012. Combined analyses of the ITS loci and the corresponding 16S rRNA genes reveal high micro- and macrodiversity of SAR11 populations in the Red Sea. PLoS ONE 7:e50274.

51. Kettler GC, Martiny AC, Huang K, Zucker J, Coleman ML, Rodrigue S, Chen F, Lapidus A, Ferriera S, Johnson J, Steglich C, Church GM, Richardson P, Chisholm SW. 2007. Patterns and implications of gene gain and loss in the evolution of Prochlorococcus. PLoS Genet 3:e231.

52. Biller SJ, Berube PM, Berta-Thompson JW, Kelly L, Roggensack SE, Awad L, Roache-Johnson KH, Ding H, Giovannoni SJ, Rocap G, Moore LR, Chisholm SW. 2014. Genomes of diverse isolates of the marine cyanobacterium Prochlorococcus. Scientific Data 1.

53. Biller SJ, Berube PM, Lindell D, Chisholm SW. 2015. Prochlorococcus: the structure and function of collective diversity. Nat Rev Microbiol 13:13–27.

54. Rocap G, Distel DL, Waterbury JB, Chisholm SW. 2002. Resolution of Prochlorococcus and Synechococcus ecotypes by using 16S-23S ribosomal DNA internal transcribed spacer sequences. Appl Environ Microbiol 68:1180–1191.

55. Ngugi DK, Antunes A, Brune A, Stingl U. 2012. Biogeography of pelagic bacterioplankton across an antagonistic temperature-salinity gradient in the Red Sea. Mol Ecol 21:388–405.

56. Somero GN, Lockwood BL, Tomanek L. 2016. Biochemical Adaptation: Response to Environmental Challenges, from Life’s Origins to the Anthropocene.

57. Haroon MF, Thompson LR, Parks DH, Hugenholtz P, Stingl U. 2016. A catalogue of 136 microbial draft genomes from Red Sea metagenomes. Scientific Data 3:160050.

58. Delmont TO, Eren AM. 2018. Linking pangenomes and metagenomes: the Prochlorococcus metapangenome. PeerJ 6:e4320.

59. Hickey DA, Singer GA. 2004. Genomic and proteomic adaptations to growth at high temperature. Genome Biol 5:117.

